# Functional characterization of two glycosyltransferases from *Withania somnifera* illuminates their role in withanosides biosynthesis and defence against bacteria

**DOI:** 10.1101/2024.02.12.579880

**Authors:** P Anjali, Ananth Krishna Narayanan, Durgesh Parihar, Anusha Patil, Dinesh A. Nagegowda

**Affiliations:** Molecular Plant Biology and Biotechnology Lab, CSIR-Central Institute of Medicinal and Aromatic Plants, Research Centre, Bengaluru - 560065, India; Academy of Scientific and Innovative Research (AcSIR), Ghaziabad-201002, India

**Keywords:** *Withania somnifera*, glycosyltransferases, biosynthesis, withanolides, withanosides, defence

## Abstract

The medicinal properties of Ashwagandha (*Withania somnifera* L. Dunal) are attributed to the presence of unique class of natural products called as withanolides and their glycosylated forms, withanosides. Withanosides are proposed to be formed from withanolides by the action of glycosyltransferases (GTs). This study reports the functional characterization of two GTs (*WsGT4* and *WsGT6*) from *W. somnifera* that exhibited induced expression in response to methyl jasmonate treatment and showed highest expression in leaves compared to other tissues. Biochemical assays with recombinant WsGT proteins showed that WsGT4 and WsGT6 formed glycosylated products with four and one of the seven tested withanolides substrates, respectively. WsGT4 catalyzed product formation using withanolide A, withanolide B, withanone, and 12-deoxywithastramonolide as substrates, with UDP-glucose serving as the glucose donor, while WsGT6 catalyzed the product formation only with withaferin A as substrate employing either UDP-glucose or UDP-galactose as sugar donors. Moreover, *in planta* studies through virus-induced gene silencing and transient overexpression of *WsGT4* and *WsGT6* in *W. somnifera* leaves modulated the levels of withanolides and withanosides, indicating their role in withanosides biosynthesis. Furthermore, while individual silencing of both *WsGT4* and *WsGT6* in *W. somnifera* reduced the tolerance to *Pseudomonas syringae* DC3000 growth, their overexpression enhanced the tolerance to the bacterium in *W. somnifera*. Taken together, these results shed light on the roles of WsGT4 and WsGT6 in withanoside biosynthesis and defence against model bacterial pathogen in *W. somnifera*.

## Introduction

Glycosylation is the process by which a carbohydrate is covalently attached to a target molecule, by the action of enzymes from a superfamily known as glycosyltransferases (GTs). GTs can glycosylate a wide range of compounds, including oligo/polysaccharides, proteins, lipids, terpenoids, flavonoids, alkaloids, and various small molecules (Sharma et al. 2007). In the context of secondary metabolic pathways, glycosylation is often regarded as a crucial final modification step resulting in the formation of a diverse range of natural glycosides (Tiwari et al. 2014). It serves as a mechanism to potentially lower the harmful effects of natural compounds, thereby altering their bioactivity, stability, solubility, subcellular localization, and binding properties (Chaturvedi et al. 2011). Among the 116 families under GT superfamily, family 1 (http://www.cazy.org/) with highest number of GTs associated with plants, features a distinctive element known as the “plant secondary product glycosyltransferase” (PSPG) box, positioned near the C-terminal end of the protein. Among the highly conserved amino acids found within the PSPG-box, the HCGWNS motif is regarded as crucial for the enzyme’s functionality (Gachon et al. 2005). Plant GTs typically utilize UDP-glucose as the primary activated sugar donor, although UDP-rhamnose, UDP-galactose, UDP-xylose, and UDP-glucuronic acid can also serve as activated sugars (Jones and Vogt 2001; Bowles et al. 2006). Glycosylation can take place at various functional groups, including -OH, -COOH, -NH2, -SH, and C-C groups, allowing for the attachment of multiple sugar moieties simultaneously (Pflugmacher and Sandermann 1998; Lim and Bowles 2004).

*Withania somnifera* (L.) Dunal, commonly known as Ashwagandha, Indian ginseng, or winter cherry, is a highly valued medicinal plant in the Solanaceae family. It is renowned for its numerous medicinal properties including antioxidative, anti-inflammatory, antidiabetic, antistress, hepatoprotective, immunomodulatory, anticonvulsant, and hypocholesterolemic activities (Bhattacharya et al. 2000; Pandey et al. 2014). These therapeutic properties are attributed to withanolides and their glycosylated derivatives, known as withanosides or glycowithanolides, found in roots (Matsuda et al. 2001; Zhao et al. 2002) and leaves (Jayaprakasam and Nair 2003). Withanosides, characterized by glucose units attached to C-3 or C-27 positions of withanolides, are suggested to play a crucial role in the plant’s defence against biotic and abiotic stresses (Chaturvedi et al. 2011; Misra et al. 2016). So far about 11 withanosides have been identified in *W. somnifera* that include withanosides I-XI (Matsuda et al. 2001; Zhao et al. 2002). While withanolides are formed from the intermediates of the phytosterol pathway derived via cycloartenol through the action of cytochrome P450 enzymes (CYP450s) (Shilpashree et al. 2022, 2024), withanosides, on the other hand, are formed from withanolides (Figure 1). This transformation occurs when withanolides undergo glycosylation, a process mediated by the action of GTs.

**Figure 1.**
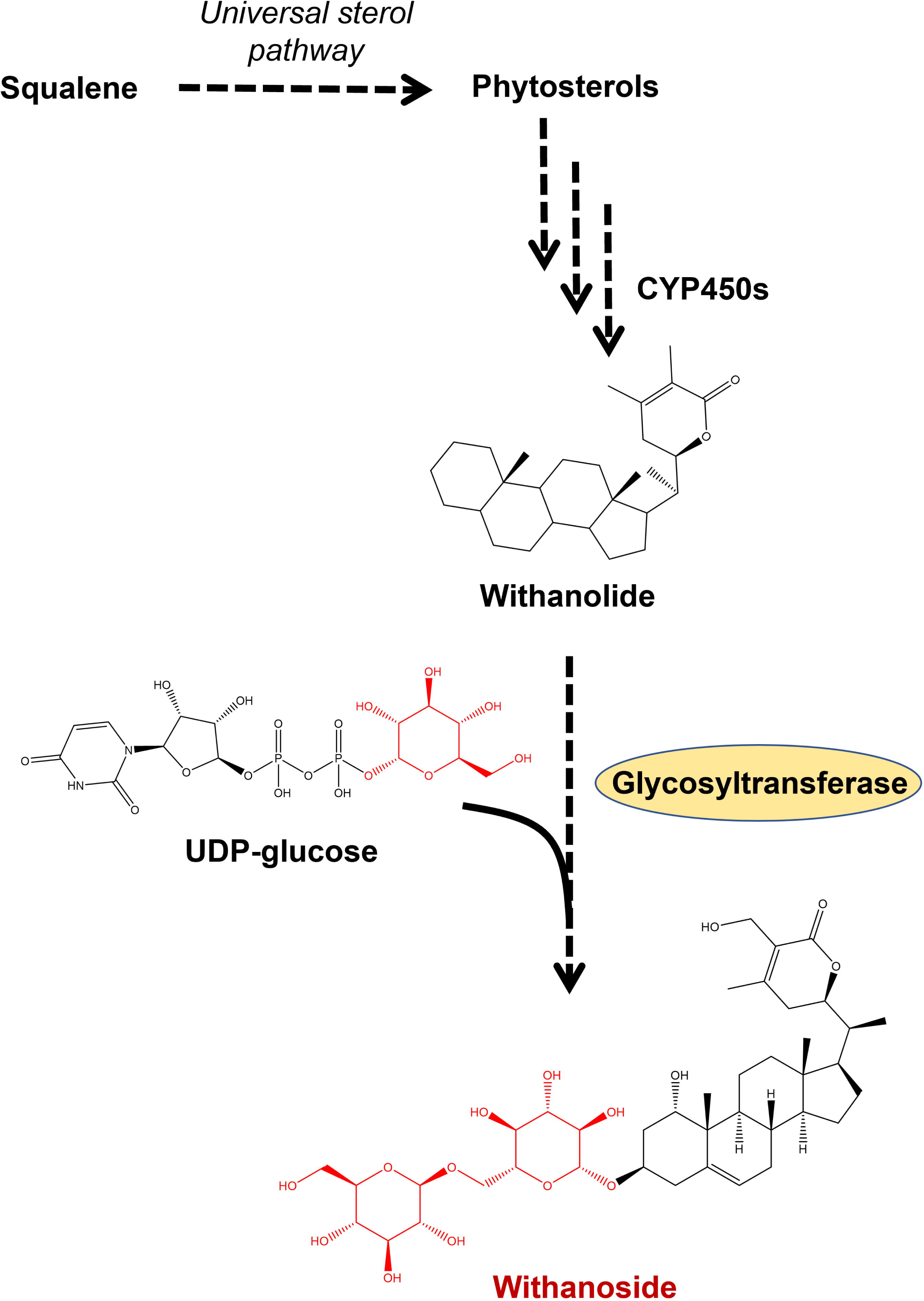
Overview of withanosides biosynthesis in *W. somnifera*. The basic structures of withanolides are decorated by sugar moieties (highlighted in red) by the action of glucosyltransferase (GT) using uridine diphosphate sugar.

Sterol glucosyltransferase WsSGTL1 was the first GT to be reported in *W. somnifera*. This *WsSGTL1* was found to be highly expressed in roots and leaves and was responsible for transferring glucose to hydroxyl group at C-3 position of sterols such as dehydroepiandrosterone, deacetyl 16-DPA, transandrosterone, 3-β-hydroxy-16, 17-α-epoxy-pregnenolene, pregnenolene, stigmasterol, β-sitosterol, ergosterol, brassicasterol, and solasodine (Sharma et al. 2007). Subsequent studies with *WsSGTL1* in transgenic lines of *W. somnifera*, *Arabidopsis thaliana*, and *Nicotiana tabacum* demonstrated its involvement in both abiotic and biotic stresses. The transgenic lines also exhibited increased levels of glycowithanolides and glycosylated sterols in *W. somnifera*, as well as glycosylated sterols and glycosylated phenolics in *A. thaliana* and *N. tabacum* (Mishra et al. 2013, 2017; Pandey et al. 2014; Saema et al. 2016). Silencing the *WsSGTL1* gene in *W. somnifera* resulted in increased accumulation of withanolides and phytosterols, while reducing the content of glycowithanolides, along with increased susceptibility to pathogens (Saema et al. 2015; Singh et al. 2016). In addition to WsSGTL1, another sterol glucosyltransferase (SGTL) named *WssgtL3.1* exhibited a protective function against salt stress and biotic stress when overexpressed in transgenic *Arabidopsis* plants (Mishra et al. 2021, 2022). Similarly, the overexpression of *Ws-Sgtl4* resulted in elevated total withanolide yield and increased levels of withanolide-A content in the transformed *W. somnifera* hairy roots (Pandey et al. 2016). Besides sterol glucosyltransferases, a GT gene encoding a flavonoid glycosyltransferase that catalyzed the regiospecific glycosylation of flavonoid-7-ols has been reported in *W. somnifera* (Kumar et al. 2013).

Despite the above studies, the enzymatic role of GTs in glycosylating withanolides and their *in planta* role in withanosides biosynthesis is poorly investigated. In this study, we have functionally characterized two GTs derived from *W. somnifera*, namely WsGT4 and WsGT6, through biochemical assays, *in planta* silencing and overexpression studies coupled with transcript and metabolite analyses, along with bacterial growth assay. The results shed light on the functions of WsGT4 and WsGT6 in withanosides biosynthesis and in bacterial defence in *W. somnifera*.

## Material and Methods

### Plant material, elicitor treatment, and tissue collection

The seeds of *W. somnifera* cv. Poshita (National Gene Bank, CSIR-CIMAP, India) were sown by casting method in a seedbed tray of 360×280×75 (l×b×h mm) dimension. The planting mixture was composed of sterile soil-rite, vermicompost, and soil in a ratio of 1:1:1. The two-leaf-staged seedlings were transplanted into pots and were maintained in a growth chamber at 25 °C with 16-hour light and 8-hour dark cycle and 70% humidity. The plants of the appropriate leaf stage were used to carry out VIGS and overexpression experiments. For methyl jasmonate (MeJA) treatment, the third developmental stage leaves were harvested from 10-week-old plants and treated with 500 µM of MeJA in a Petri dish. The tissues were collected at different time (h) intervals and stored at -80°C until further use. For tissue-specific gene expression analysis, berries, flowers, leaves, roots, and stems were collected from 10-week-old *W. somnifera* plants and stored at -80 °C for further use (Singh et al. 2015, 2017).

### Phylogenetic analysis

The sequences of *WsGT4* and *WsGT6* were mined using our *W. somnifera* in-house leaf transcriptome. The Open Reading Frame (ORF) of *WsGT4* (1356 bp) and *WsGT6* (1419 bp) was detected using the NCBI ORF Finder (https://www.ncbi.nlm.nih.gov/orffinder/). The molecular weight of WsGT4 (451 Aa, 51005.11 kDa) and WsGT6 (472 Aa, 53057.75 kDa) was predicted on ExPASy. The conserved domains were analysed in NCBI Conserved Domain Database (https://www.ncbi.nlm.nih.gov/Structure/cdd/wrpsb.cgi). A phylogenetic tree was constructed by the neighbor-joining method using the MEGA 11 software with 1000 bootstrap replicates. Multiple sequence alignment with closely related GTs from other plant species was performed using ESPript 3.0 (Robert and Gouet 2014).

### RNA isolation, cDNA synthesis and qRT-PCR analysis

RNA was isolated, cDNA was prepared, and qRT-PCR analysis was conducted as previously described in the literature (Singh et al. 2015, 2017). The cDNA was normalized using *18S* rRNA as an internal reference. Gene-specific primers, designed outside the gene region utilized for cloning into pTRV2 were employed for the qRT-PCR analysis (Table S1). The reaction was carried out using 2X SYBR green mix from Thermo Scientific (USA) and was run in the StepOne Real-Time PCR System from Applied Biosystems (USA). The qRT-PCR conditions used were as follows: 94 °C for 10 min, followed by 40 cycles of 94 °C for 15s and 60 °C for 45 sec. Fold-change differences in gene expression were analyzed using the comparative cycle threshold (*C*_t_) method.

### Glycosyltransferase enzyme assay

The open reading frame of *WsGT4* and *WsGT6* was cloned into the *Nhe*I/*Hind*III and *Nde*I/*Bam*HI sites of pET28a bacterial expression vector in-frame with the C-terminal 6X-His-tag (Novagen, http://www.emdbiosciences.com) resulting in pET28a::*WsGT4* and pET28a::*WsGT6* respectively (Figure S1 A&B). For recombinant protein expression, pET28a::*WsGT4*, pET28a::*WsGT6* and pET28a empty vector (control) constructs were transformed into *E. coli* BL21 Rosetta-2 cells. Expression of recombinant proteins were induced when OD_600_ of the culture was 0.4 by the addition of 0.1 mM isopropylthio-β-galactoside (IPTG) and grown in an incubator shaker at 18 °C, 180 rpm for 16 hrs. The recombinant protein was purified by affinity chromatography using nickel-nitrilotriacetic acid (Ni-NTA) agarose beads (Qiagen, http://www.qiagen.com) as per the manufacturer’s instructions. The concentration of the purified protein was determined using the Bradford method (Bradford, 1976) and purity was determined using SDS-PAGE analysis (Figure S1 C&D). Enzyme assays were performed using purified recombinant proteins of WsGT4 and WsGT6. The assay buffer comprised of 10 mM Tris-HCl of pH 7.5 and 1 mM DTT. Two µg each of purified WsGT4 and WsGT6 was incubated with 1 mM of UDP-glucose/galactose and 50 µM of withanolide substrates in a 500 µl reaction for 16 h at 30°C. The reaction was stopped by the addition of equal volumes of a 7:3 chloroform: methanol solution. The metabolites were extracted twice using the 7:3 chloroform: methanol solution, pooled and dried using a vacuum concentrator (Eppendorf). The dried samples were resuspended in 100 µl of methanol and subjected to HPLC analysis.

### Generation of Virus-Induced Gene Silencing (VIGS) and overexpression constructs

Plasmids pTRV1 and pTRV2 (Liu et al. 2002) procured from TAIR (www.arabidopsis.org) were used to generate VIGS constructs. A 470-bp fragment corresponding to *WsGT4* and a 492-bp fragment corresponding to *WsGT6* were amplified from leaf cDNA using gene-specific primers (Table S1). Care was taken to select the region that is specific for each gene to avoid off-target silencing (Figure S2). The amplicons were cloned into pJET1.2/blunt vector for sequence confirmation. Further, *WsGT4* and *WsGT6* were subcloned into *Xba*I and *Xho*I, and *Xba*I and *Bam*HI sites of pTRV2 vector yielding pTRV2::*WsGT4* and pTRV2::*WsGT6* constructs, respectively (Figure S3A). For the generation of overexpression constructs, the ORF of *WsGT4 and WsGT6* was PCR amplified using leaf cDNA with specific forward and reverse primers (Table S1). The amplicons were cloned into pJET1.2/blunt vector and confirmed by nucleotide sequencing. Later, *WsGT4* and *WsGT6* fragments were subcloned into the *Eco*RI and *Sac*I, and *Nde*I and *Bam*HI sites of the pRI101-AN binary vector under the control of the *35S* promoter of Cauliflower mosaic virus (CaMV) to form the pRI101-AN::*WsGT4* and pRI101-AN::*WsGT6* constructs, respectively (Figure S3B). The generated VIGS and overexpression constructs were introduced into GV3101 strain of *Agrobacterium tumefaciens* by the freeze–thaw method.

### VIGS and transient overexpression of *WsGT*s

Virus-induced gene silencing experiments were carried out by infiltration of leaves in *W. somnifera* plants according to Singh et al., 2015. Briefly, the overnight grown cultures of *Agrobacterium* were pelleted and resuspended in infiltration buffer composed of 10 mM MgCl_2_, 10 mM MES and 100 µM acetosyringone, pH 5.6. The OD_600_ was adjusted to 1.3 and incubated for 4-5 hours at 28°C. Infiltration was performed using 1:1 ratio of *Agrobacteria* cultures harboring TRV1 and TRV2 or its derivatives using 1-mL needleless syringe on the abaxial side of leaves in 4-leaf-staged plants. Thirty days-post-infiltration (dpi), tissues exhibiting viral infection phenotype were collected for transcript and metabolite analyses. For transient overexpression, overnight grown *Agrobacterium* cultures were pelleted by centrifugation and resuspended in infiltration buffer (50 mM MES, 2 mM Na_3_PO_4_, 0.5% glucose, and 100 µM acetosyringone with pH adjusted to 5.6). The OD_600_ was adjusted to 0.2 and incubated at 28°C for 3-4 hours. Leaf infiltration was performed as described above into the first pair of leaves of 6-8 leaf-staged plants and placed in dark condition for 48 hours. The infiltrated leaves were collected and stored at -80°C for further metabolite and transcript analysis (Singh et al. 2015).

### Extraction and analysis of withanolides and withanosides

Withanolides and withanosides extraction and analysis was performed following Singh et al., 2015. Briefly, 20 mg of dried leaf tissue was ground using 1 mL absolute methanol followed by sonication. The supernatant was collected, and the ground tissue was extracted again in 1 mL methanol followed by subsequent sonication, this step was repeated. Methanolic leaf extract was collected in a scintillation vial and allowed to evaporate, and the residue was resuspended in 3 mL of 70% methanol. This was extracted thrice with 3 mL chloroform. The lower layer of chloroform extract (9 mL) was decolorized using charcoal followed by drying. The dried samples were treated with 200 µL of chloroform: methanol in the ratio of 7:3 and used for HPLC analysis. The HPLC (Model: Nexera, Shimadzu, Kyoto, Japan) fitted with C-18 (250 mm x 4.60 mm, 5 µm) column was used for metabolite analysis. The program used has been previously described in Singh et al., 2015. Withanolide and withanoside standards were procured from Natural Remedies Pvt. Ltd. (Bangalore, India). The quantity of sample used for injection was 20 µL. Catharanthine was used as an internal standard. The area of individual withanolides and withanosides was determined after normalizing with the peak area of the internal standard, catharanthine (Sigma-Aldrich) (Singh et al. 2015).

### Bacterial growth curve assay

The model plant pathogen *Pseudomonas syringae* pv. tomato DC3000 was inoculated in 5 mL nutrient broth and grown overnight in an incubator shaker at 28°C. The culture was centrifuged and the bacterial pellet was resuspended in 10 mM MgCl_2_ and the cell density was adjusted to OD_600_=0.002. Bacterial infiltration and colony forming units (cfu) determination were performed according to Singh et al. 2015, 2017. To assess the bacterial infection rate, leaf discs from infected and buffer infiltrated control plants were collected from the zone of infiltration 3 days post inoculation. Leaf discs were homogenized using 10 mM MgCl_2_ and the cfu/cm^2^ was determined by plating serial dilutions of leaf extracts on *Pseudomonas*-specific agar plates (Singh et al. 2015, 2017).

### Statistical analysis

Mean, standard deviation and standard error was calculated using GraphPad QUICKCALC online tool. The statistical significance between control and treated samples was calculated using unpaired Students t-test. Asterisks indicate significant difference between samples (\**p*<0.05; \*\**p*<0.01; and \*\*\**p*<0.001).

## Results

### Identification of WsGTs and sequence analysis

The sequences corresponding to GTs were mined and retrieved from the in-house generated leaf transcriptome of *W. somnifera*. Two *GT* candidates with complete ORF that were highly induced in response to MeJA were considered for further characterization. *WsGT4* comprised an ORF of 1356 bp, encoding a protein of 451 amino acids with a calculated molecular mass of 51.0 kDa (accession number: WsGT74E2). On the other hand, *WsGT6* contained an ORF of 1419 bp, encoding a protein of 472 amino acids with a calculated molecular mass of 53.06 kDa (accession number: WsGT75C1). Phylogenetic analysis of WsGT4 and WsGT6 with other reported GTs from *W. somnifera* and GTs of other plant species indicated that WsGT4 has 75.78% identity with UGT73A10 from *Lycium barbarum* followed by 73.54% identity with NtUGT103 from *Nicotiana tabacum*, whereas WsGT6 exhibited 86.29% identity with SlUGT75C1 from *Solanum lycopersicum* followed by 85.59% identity with NtGT2 from *Nicotiana tabacum* (Figure 2A). Multiple sequence alignment of WsGT4 and WsGT6 with closely related GTs from other plant species showed that they belonged to family 1 of plant GT superfamily that has the conserved HCGWNS motif within PSPG box, a characteristic feature of GTs involved in secondary metabolism (Figure 2B).

**Figure 2:**
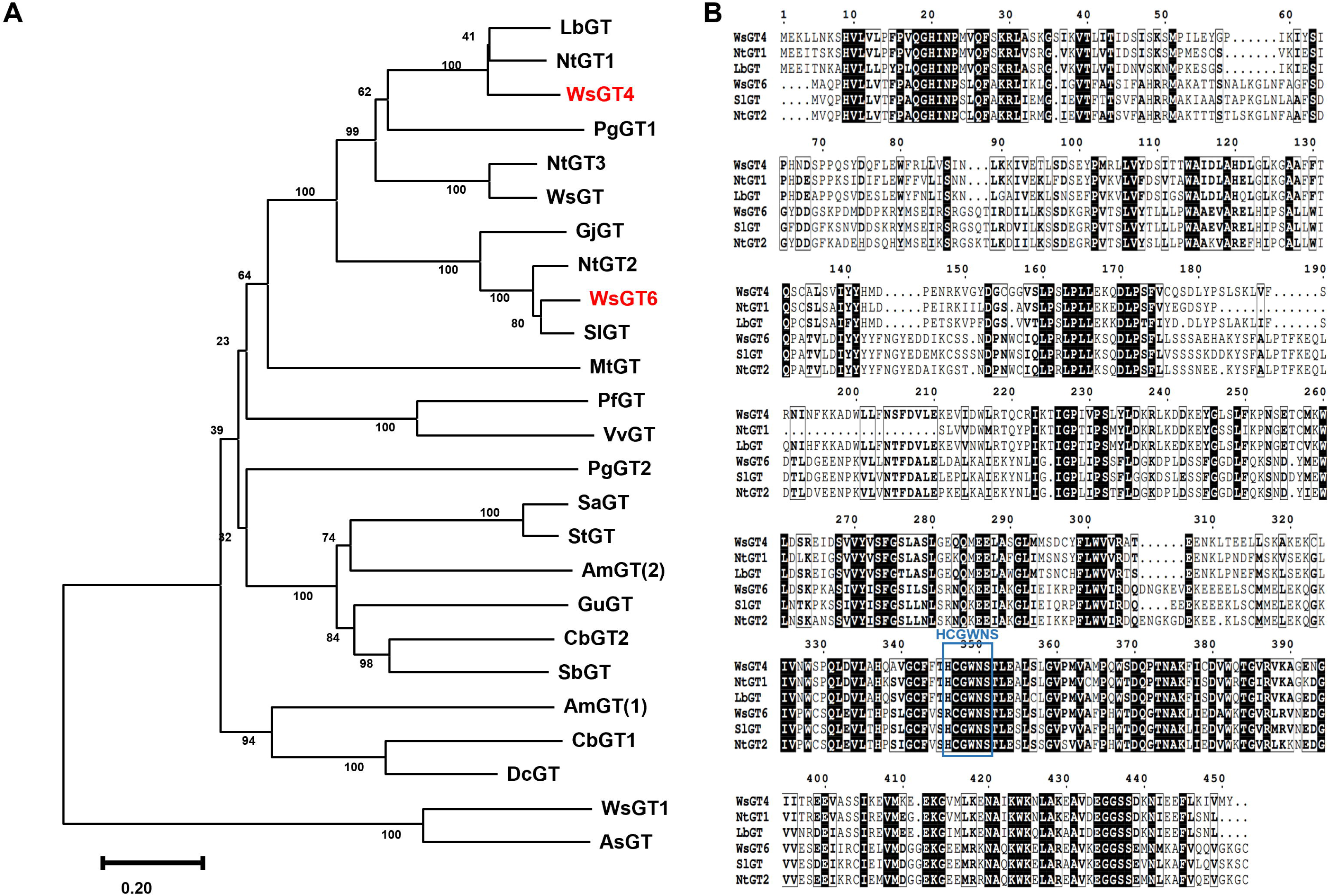
Sequence analysis of WsGT4 and WsGT6 with other related plant GTs. **(A**) Phylogenetic tree constructed by the neighbour-joining method using MEGA 11, showing sequence relatedness of WsGT4 and WsGT6 with other GTs, Lb-*Lycium barbarum*, Nt-*Nicotiana tabacum*, Ws-*Withania somnifera*, Pg-*Panax ginseng*, Gj*-Gardenia jasminoides*, Sl*-Solanum lycopersicum*, Mt-*Medicago truncatula*, Pf-*Perilla frutescens*, Vv-*Vitis vinifera*, Sa-*Solanum aculeatissimum*, St-*Solanum tuberosum*, Am(2)-*Astragalus membranaceus*, Gu-*Glycyrrhiza uralensis*, *Cb-Cleretum bellidiforme*, Sb-*Scutellaria baicalensis*, Am(1)-*Antirrhinum majus*, Dc-*Dianthus caryophyllus*, As-*Avena sativa*. **(B)** Multiple Sequence Alignment of WsGT4, WsGT6, NtGT1, NtGT2, LbGT, and SlGT (performed using ESPript 3.0) showing the presence of HCGWNS motif, Ws-*Withania somnifera*, Nt-*Nicotiana tabacum*, Lb-*Lycium barbarum*, and Sl*-Solanum lycopersicum*. NCBI Accessions: WsGT4: WsGT74E2, WsGT6: WsGT75C1, LbGT: BAG80543.1, NtGT1: WIW42716.1, NtGT3: AAF61647.1, WsGT: ACM09899.2, PgGT1: AGR44631.1, GjGT: BAM28983.1, NtGT2: BAB88935.1, SlGT: NP_001348274.1, MtGT: pdb|2PQ6|A, PfGT: BAA19659.1, VvGT: AAB81682.1, PgGT2: AGR44632.1, SaGT: BAD89042.1, StGT: ABB29874.1, AmGT(2): UIW25028.1, GuGT: AXS75258.1, CbGT2: CAB56231.1, SbGT: BAA83484.1, AmGT(1): BAE48239.1, CbGT1: AAL57240.1, DcGT: BAD52004.1, WsGT1: ABC96116.1, AsGT: CAB06081.1

### Spatio-temporal expression of *WsGT4* and *WsGT6*

MeJA is a well-known inducer of pathway genes involved in secondary metabolism and plant defence response, including genes related to withanolide biosynthesis (Shilpashree et al. 2022). To determine whether expression of *WsGT4* and *WsGT6* were altered in response MeJA, qPCR was performed. It was found that the expression of *WsGT4* drastically increased after 6 and 48 hours of MeJA treatment, whereas *WsGT6* exhibited a drastic induced expression only after 48 hours upon exposure to MeJA (Figure S4). The observed MeJA-induced expression of *WsGT4* and *WsGT6* indicated their possible involvement in secondary metabolism, plant defence and withanosides biosynthesis. Further, tissue-specific expression analysis revealed that *WsGT4* and *WsGT6* had the highest expression in leaves compared to other tissues. Both genes exhibited a similar expression pattern across different tissues, with leaf showing the highest expression, followed by root, flower, and berry, while stem exhibited the lowest expression levels. Among the two genes, *WsGT6* exhibited higher overall expression levels in all tissues compared to *WsGT4*. Both *WsGT4* and *WsGT6* showed highest expression in leaf with ∼28-fold and ∼100-fold increased expression compared to the stem of *WsGT4* sample (which was set to 1), respectively. In terms of expression between the two genes, *WsGT6* displayed ∼4 times higher expression in leaves and about 5-10 times higher expression in other tissues compared to *WsGT4* (Figure 3).

**Figure 3:**
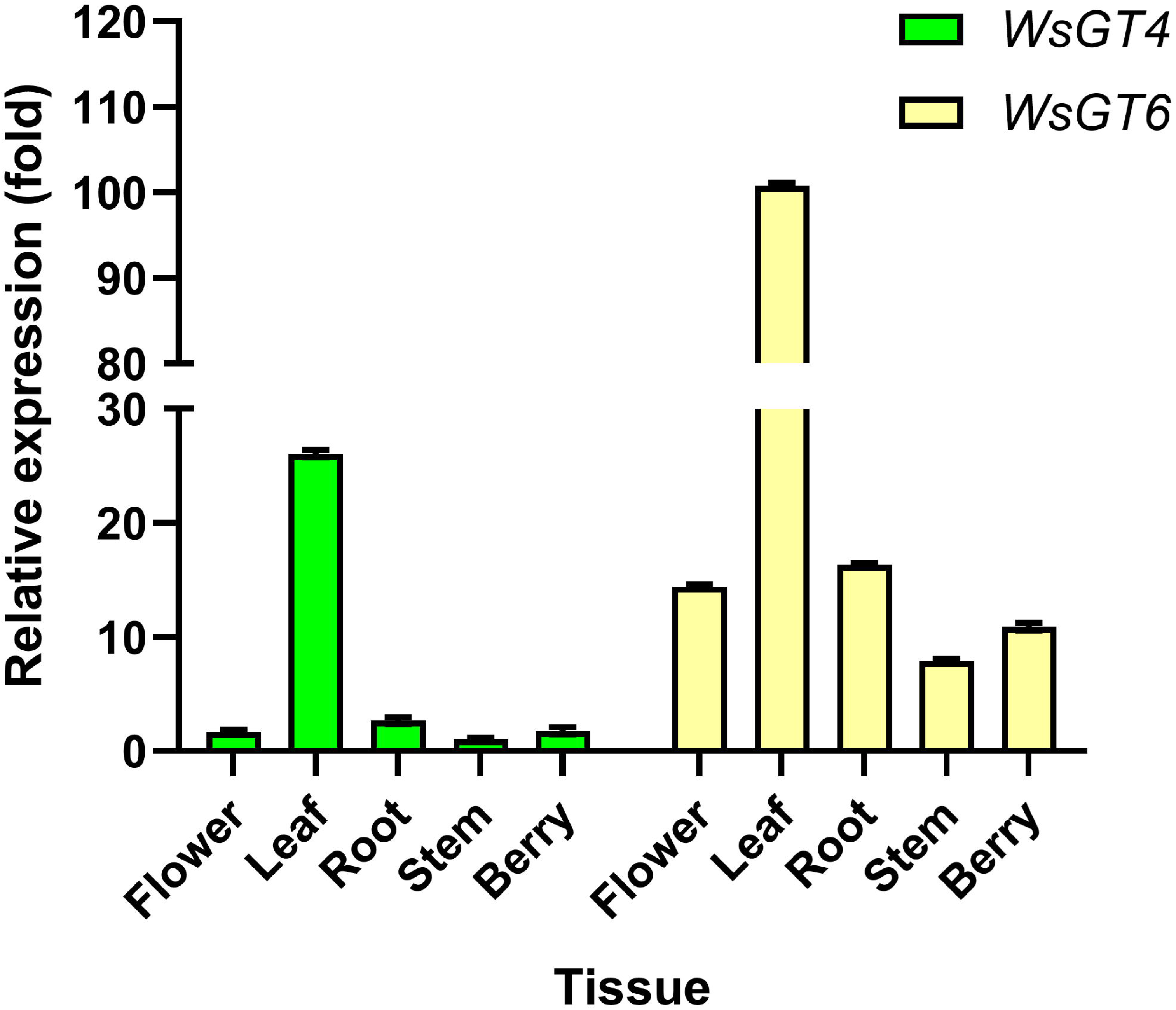
Tissue specefic expression of *WsGT4* and *WsGT6*. qRT-PCR analysis was performed to check the expression of *WsGT4* (green) and *WsGT6* (yellow) in the leaf, flower, root, stem and berry of *W. sominifera*. The data is represented as fold change in expression with respect to the expresstion of *WsGT4* in the stem which was set to 1. It was found that both genes are maximally expressed in the leaf. Error bars represent SEM and **** represents a *p*<0.0001.

### Biochemical characterization of WsGT4 and WsGT6

Enzyme assays were performed to determine the biochemical function of WsGT4 and WsGT6. Recombinant proteins WsGT4 and WsGT6 were expressed in *E. coli* and purified using Ni-NTA affinity chromatography. These purified proteins were incubated with combinations of seven different withanolides substrates along with either UDP-glucose or UDP-galactose as a sugar donor. Analysis of the reaction products using HPLC revealed that among the tested substrates, WsGT4 catalyzed product formation when incubated with withanolide A, withanolide B, withanone, and 12-deoxywithastramonolide in the presence of UDP-glucose as the sugar donor (Figure 4) but, there was no product formation with withanoside IV, withanoside V, and withaferin A (Figure S5). In contrast, no product was formed with any of the tested withanolides substrates when UDP-galactose was used as a sugar donor (Figure S6). Furthermore, WsGT4 catalyzed the formation of two distinct products with different retention times when withanolide A, withanolide B, and withanone were used as substrates, whereas it formed only one product in the presence of 12-deoxywithastramonolide. The efficiency of product formation by WsGT4 was higher with withanolide A, withanolide B, and withanone substrates compared to 12-deoxywithastramonolide substrate. Contrary to the glycosyltransferase activity of WsGT4, assays using WsGT6 showed that it catalyzed the product formation only when incubated with withaferin A and either UDP-glucose or UDP-galactose as the sugar donor. Though WsGT6 produced five different products with varying retention times, it exhibited a similar pattern of product formation in the presence of both UDP-glucose and UDP-galactose as the sugar donor (Figure 5). WsGT6 did not show any product formation with withanolide A, withanolide B, withanone, 12-deoxywithastramonolide, withanoside IV, and withanoside V substrates and either UDP-glucose or UDP-galactose as the sugar donor (Figure S7 & S8). Though HPLC analysis showed WsGT4 and WsGT6 formed glycosylated products in presence of withanolides substrates, the exact chemical nature of the product formed could not be determined by LC-MS due to similarities in the fragmentation patterns of the substrate and products.

**Figure 4:**
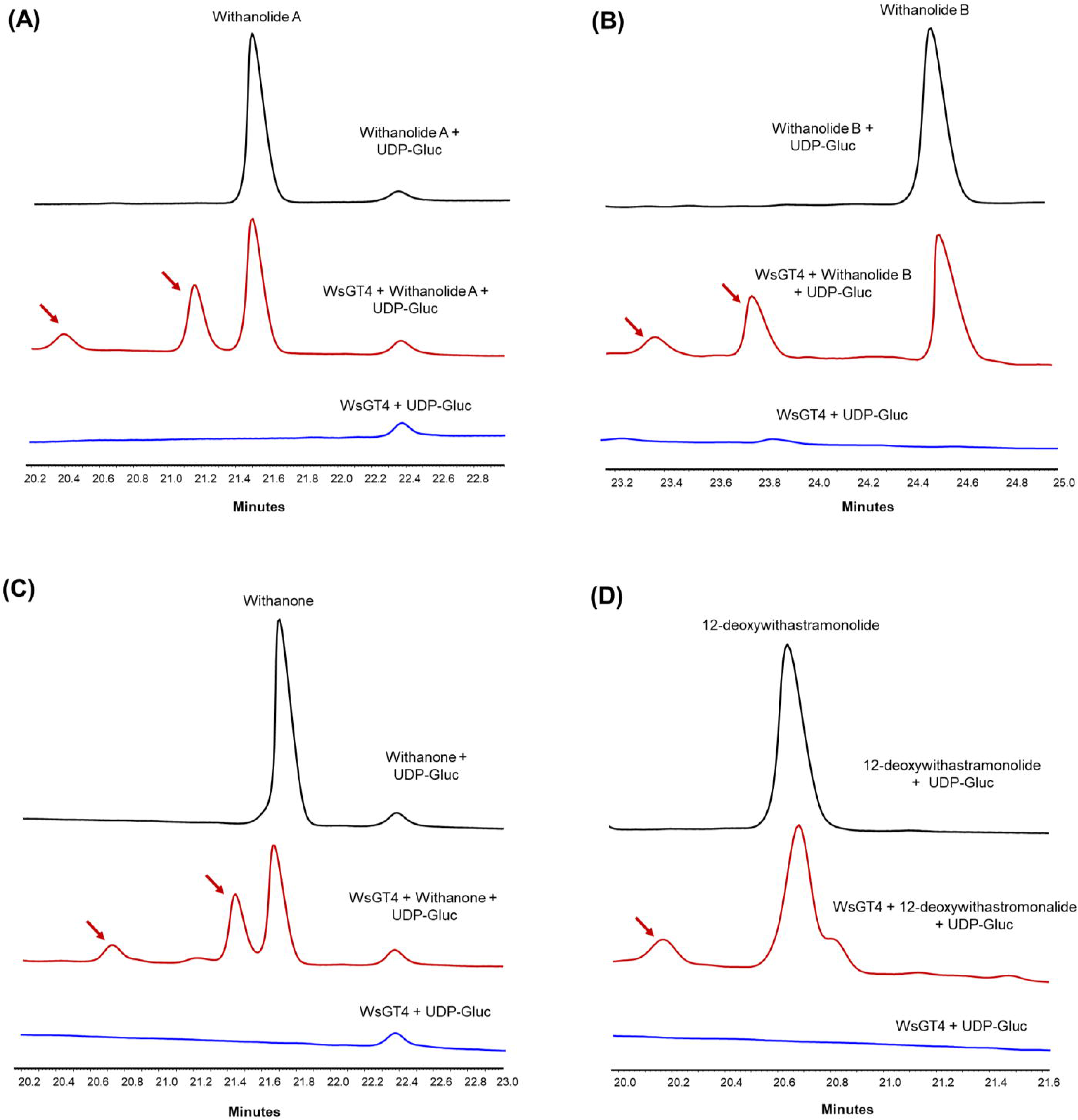
WsGT4 exhibits glycosyltransferase activity with different withanolides. Purified recombinant WsGT4 protein was incubated with different withanlides and withanosides along with UDP-glucose as a sugar donor. Analysis of reaction products by HPLC showed product formation by WsGT4 in presence of withanolide A **(A)**, withanolide B **(B)**, withanone **(C)** and 12-deoxywithastramonolide **(D)** as substrates, with UDP-glucose. Arrows indicate the product peak formed in the assay.

**Figure 5:**
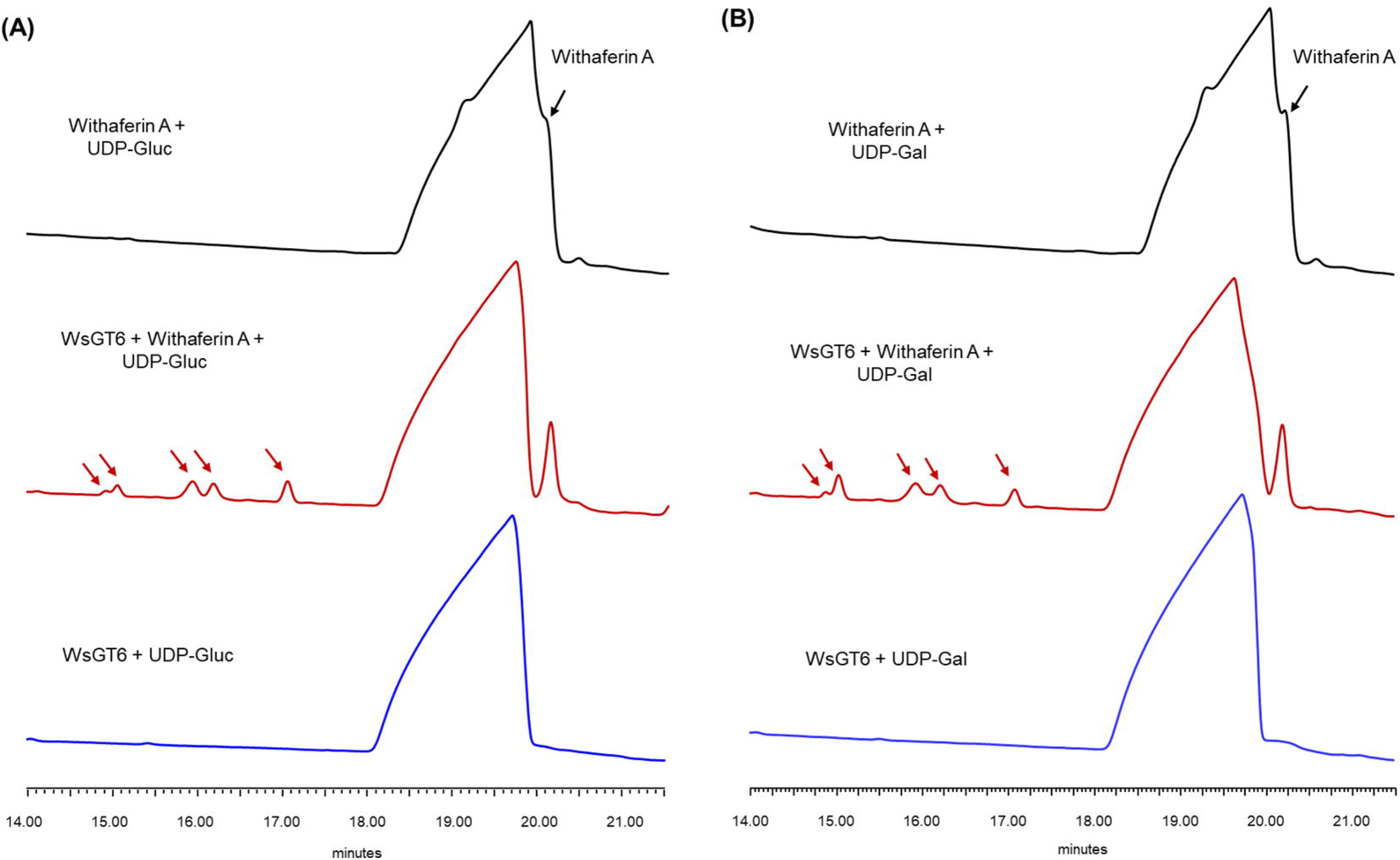
Glycosyltransferase assay for WsGT6 with withaferin A as substrate. Purified recombinant WsGT6 protein was incubated with withaferin A along with UDP-glucose (UDP-Glc) or UDP-galactose (UDP-Gal) as sugar donors. Analysis of reaction products by HPLC showed product formation by WsGT6 in presence of withaferin A as substrate with UDP-glucose **(A)** or UDP-galactose **(B)**. Arrows indicate the product peak formed in the assay.

### Effect of *WsGT4* and *WsGT6* silencing on withanolide and withanoside accumulation

To determine the *in planta* role of *WsGT4* and *WsGT6* in withanolide and withanoside biosynthesis, VIGS was performed. As VIGS is prone to off target silencing (Dinesh-Kumar et al.), the sequence region unique to *WsGT4* and *WsGT6* was chosen to generate the silencing constructs. Thirty dpi, leaves of similar developmental stages exhibiting typical viral infection symptoms were collected from *WsGT4*-VIGS and *WsGT6-*VIGS plants along with empty vector (control) plants for transcript and metabolite analysis. qRT-PCR analysis revealed that the expression of *WsGT4* and *WsGT6* was down-regulated by about 90% and 88%, respectively, in *WsGT4*-VIGS and *WsGT6-*VIGS plants compared to the control plant (Figure 6B&D). Subsequent metabolite analysis by HPLC in *WsGT4*- and *WsGT6*-silenced, as well as control plants, revealed a differential modulation in the levels of various withanolides and withanosides in silenced plants. The silencing of *WsGT4* resulted in significantly elevated levels of withanoside IV (194.8%), withanolide A (189.6%) and withanolide B (235.4%), whereas silencing of *WsGT6* resulted in elevated levels of 12-deoxywithastramonolide (256.3%) and withanolide A (285.26%) when compared to unsilenced plants (Figure 6C&E).

**Figure 6:**
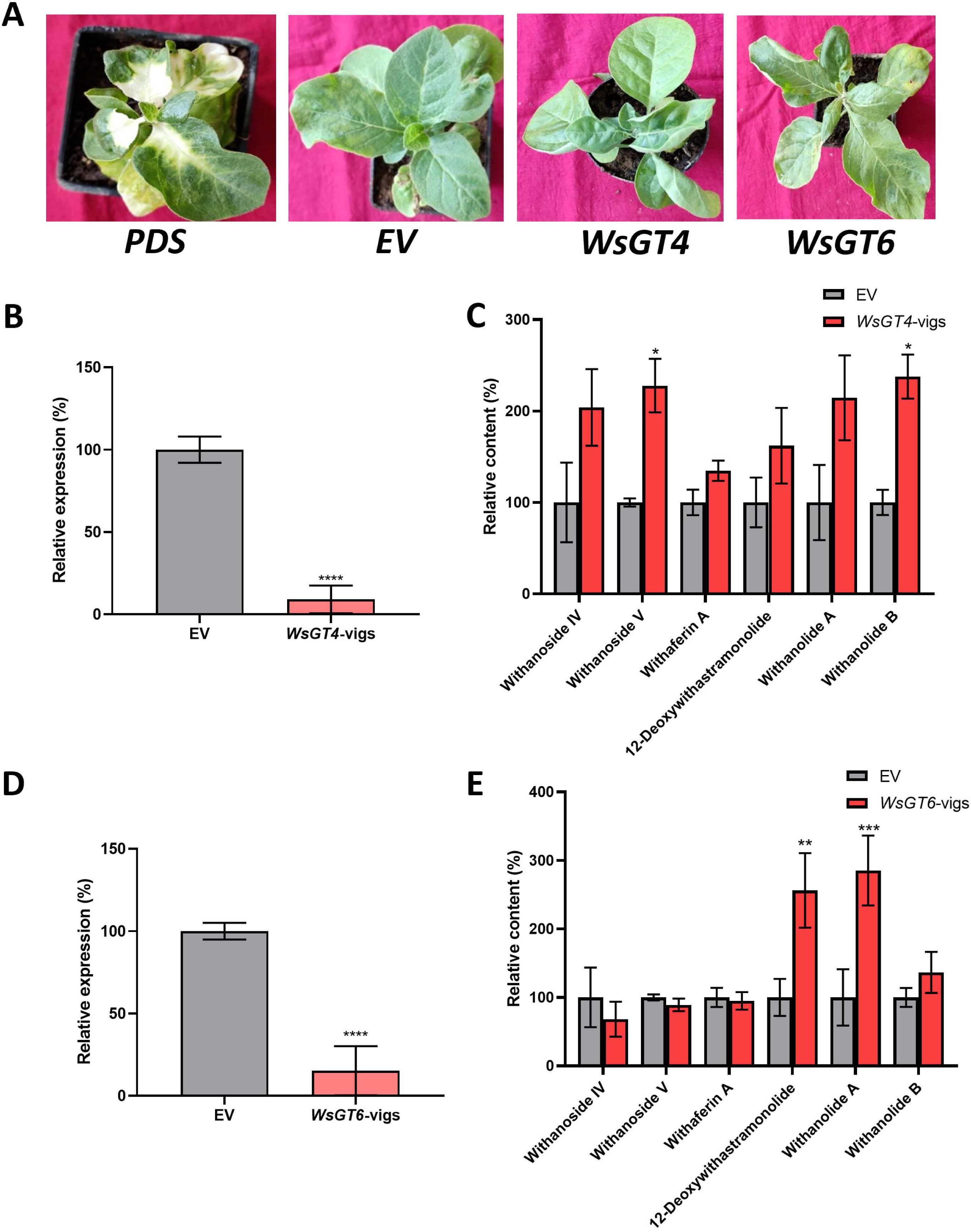
Effect of *WsGT4-* and *WsGT6-* silencing on withanolides content in *W. somnifera*. **(A)** Phenotype of *W. somnifera* plants infiltrated with pTRV2 empty vector control (EV), pTRV2-*PDS*, pTRV2-*WsGT4,* and pTRV2-*WsGT6* constructs. **(B and D)** Relative gene expression of *WsGT4* **(B)** and *WsGT6* **(D)** measured in silenced leaves of *WsGT4*-VIGS and *WsGT6*-VIGS samples in comparison to EV. The expression levels of *WsGT4* and *WsGT6* transcript were normalized to *18S rRNA*. Data are represented as percentage relative expression with the expression of genes in the EV plant set as 100%. Error bars represent SEM of three biological replicates, each having three technical replicates. **(C and E)** HPLC quantification of witanolides and withanosides in silenced leaves of EV control, *WsGT4*-VIGS **(C)** and *WsGT6*-VIGS **(E)** samples. The levels of metabolites in the silenced leaves (red bars) are expressed in terms of relative percentage in comparison to that of EV control (grey bars) which was set at 100%. Error bars represent SEM of 4 independant experiments, * represents a *p*<0.05, ** represents *p*<0.01, *** represents *p*<0.001 and **** represents *p*<0.0001.

### Overexpression of *WsGT4* and *WsGT6* and its effect on withanolide and withanoside biosynthesis

Silencing of *WsGT4* and *WsGT6* resulted in the modulation of withanolides and withanosides in *W. somnifera*. To further explore the impact of overexpressing these genes on withanolides and withanosides biosynthesis, we conducted transient overexpression of *WsGT4* and *WsGT6* in *W. somnifera*. This was achieved by infiltrating leaves with *Agrobacteria* carrying the respective overexpression constructs. On three dpi, the infiltrated leaves were collected for metabolite and transcript analysis. Gene expression analysis revealed a substantial increase in expression levels, with *WsGT4* and *WsGT6* being overexpressed approximately 17.6-fold and 2.3-fold, respectively, compared to expression in empty vector-infiltrated samples (Figure 7A&C). Metabolite analysis of *WsGT4*-overexpressing leaves indicated a significant decrease in the levels of withanoside IV, withanoside V, and 12-deoxywithastramonolide, dropping to 22.16%, 16.48%, and 52.98% of the levels observed in control plants, respectively (Figure 7B). On the other hand, overexpression of *WsGT6* led to a notable decrease in the levels of 12-deoxywithastramonolide and withanolide A, reducing to approximately 63.95% and 47.9% of the levels in control plants, respectively. Interestingly, overexpression of *WsGT6* also resulted in a significant increase in withaferin A levels, with an enhancement of 139.2% in the leaves (Figure 7D).

**Figure 7:**
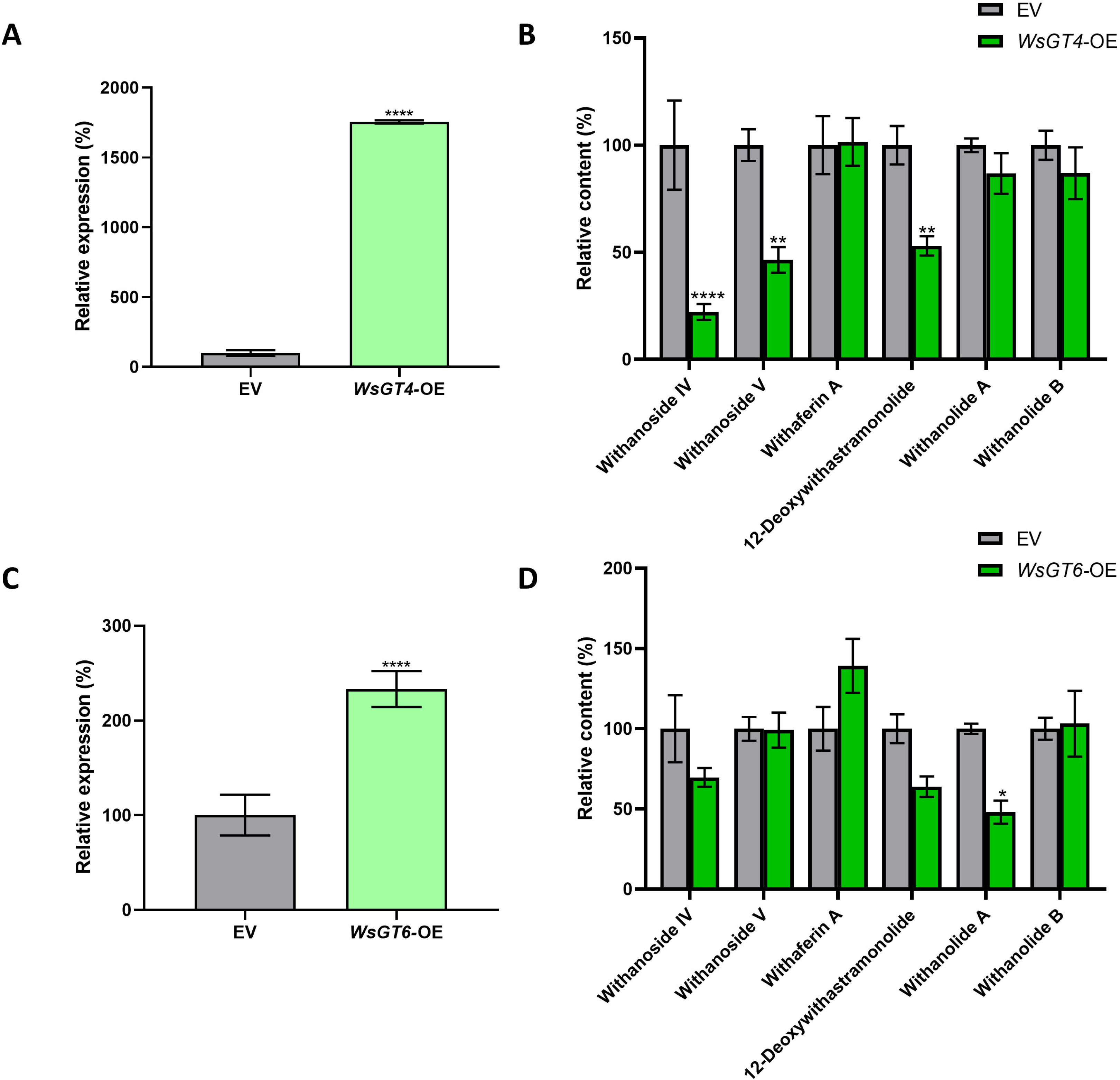
Effect of overexpression of *WsGT4* and *WsGT6* on withanolide content. Transient overexpression of *WsGT4* (**A**, *WsGT4*-OE) and *WsGT6* (**C**, *WsGT6*-OE) was performed using pRI101-AN plant overexpression vector, the expression of the overexpressed genes was analysed using qRT-PCR and compared with that of empty vector infiltrated leaves (Control). The data is represented as percentage expression with the expression of genes in the control plant set as 100%. Overexpression resulted in significant increase in expression of *WsGT4* and *WsGT6* to about 1795% and 233% of that in the control plants respectively. Leaves infiltrated with empty pRI101-AN vector (control) and overexpression construct (OE) construct of *WsGT4* **(B)** and *WsGT6* **(D)** were collected and their metabolites were analyzed using HPLC. The levels of metabolites in the overexpressed leaves (green bars) are expressed in terms of percentage in comparison to that of control (grey bars) which was set at 100%. Error bars represent SEM 9 independant experiments, * represents a *p*<0.05, ** represents *p*<0.01, *** represents *p*<0.001 and **** represents *p*<0.0001.

### Role of WsGT4 and WsGT6 in defence against bacteria

As silencing and transient overexpression of *WsGT4* and *WsGT6* resulted in modulation of withanolides and withanosides content, bacterial growth assay was performed to determine their role in defence. Leaves from *WsGT4* and *WsGT6* silencing background developed severe disease symptoms and sustained more tissue damage than control leaves (Figure S9 A). Further, bacterial growth assay using extract isolated from infiltrated leaves 3 dpi showed that both *WsGT-*VIGS samples exhibited drastic increase in the growth of *P. syringae* DC3000 than that of control. The log cfu/cm^2^ for *WsGT4-* and *WsGT6*-VIGS was found to be 6.5 cfu/cm^2^ whereas it was ∼6 cfu/cm^2^ for control (Figure 8A). In contrast to the effect of VIGS on bacterial growth, overexpression of *WsGTs* led to enhanced tolerance towards bacterial growth. Leaves overexpressing *WsGT4* and *WsGT6* remained healthy, and exhibited no necrotic phenotype compared to control at 4 dpi of *P. syringae* DC3000 (Figure S9 B). Subsequent bacterial growth assay revealed a significant reduction in the growth of *P. syringae*. The log cfu/cm^2^ in *WsGT4* and *WsGT6* overexpressing tissues was ∼4 while it was 5.2 in control (Figure 8B). Hence, it is possible that the products formed by WsGT4 and WsGT6 enzymes could be involved in providing tolerance through their antibacterial activity.

**Figure 8:**
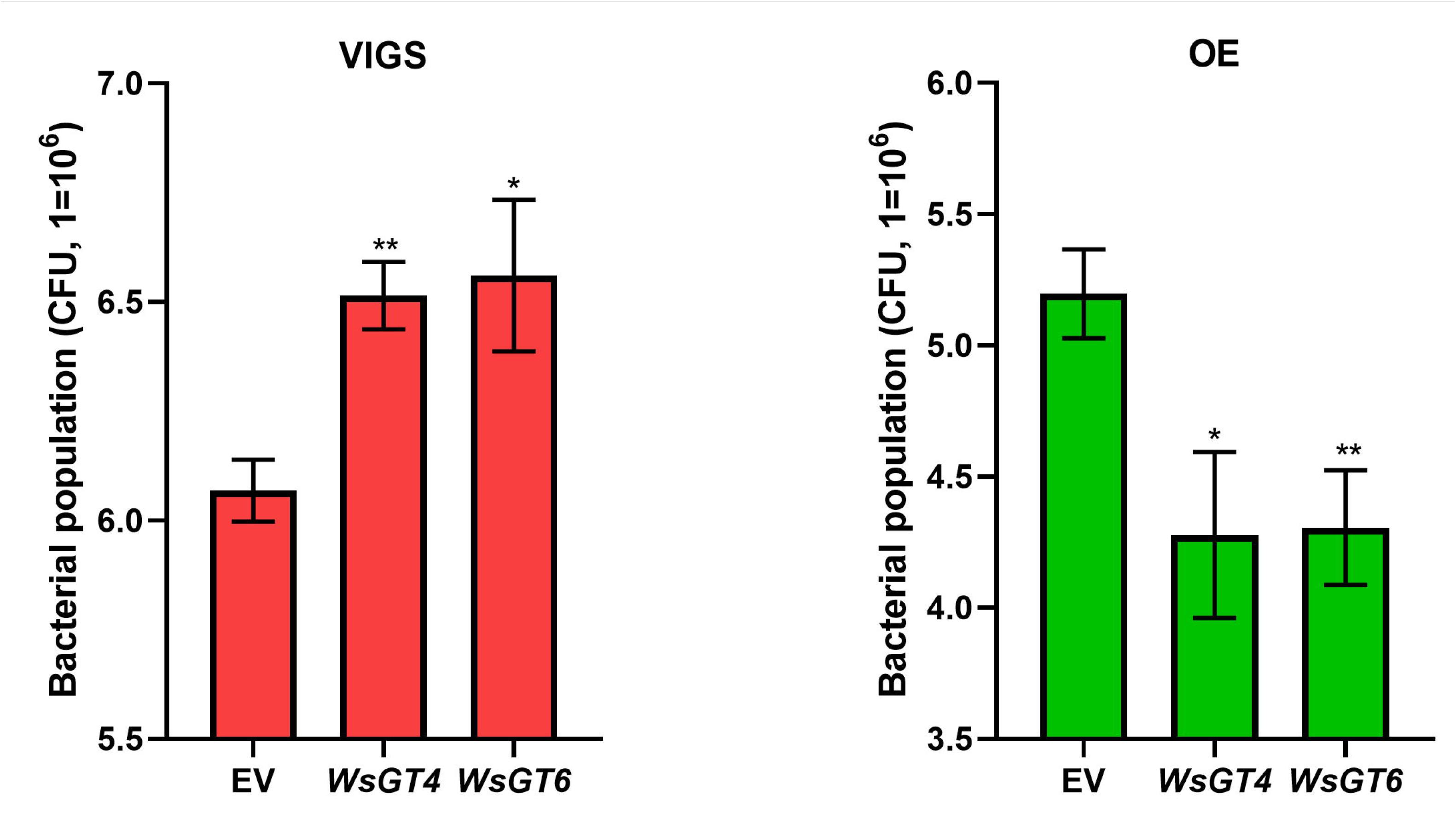
Effect of *WsGT4* and *WsGT6* expression on *Psuedomonas syringae* growth. Leaves of *W. sominifera* were infiltrated with empty vector (EV), silencing (VIGS, **A**) and overexpression (OE, **B**) constructs of *WsGT4* and *WsGT6* and simultaneously infiltrated with *P. syringae* pv. tomato DC3000 cells. 3 days post infiltration, infiltrated leaves were collected, homogenized, serially diluted, and plated on *Pseudomonas*-specific agar plates to determine the colony forming units (CFU) of *Pseudomonas syringae*. Error bars show meanLJ±LJSE of three independent experiments. Statistical significance represented as \**p*<LJ0.05, \*\**p*<LJ0.01 a significant difference.

## Discussion

Withanosides are the glycosides of withanolides, and are key contributors to the medicinal properties attributed to *W. somnifera*. They are formed by the action of GTs that transfer glycosyl group to withanolides. Despite some studies characterizing GTs in *W. somnifera* (Sharma et al. 2007; Kumar et al. 2013), none have reported GTs that are directly involved in glycosylating withanolides. In this study, we report the characterization of two new GT genes from *W. somnifera*, namely *WsGT4* and *WsGT6*. Their encoded proteins specifically glycosylate certain withanolides, modulate the *in planta* levels of both withanolides and withanosides, and provide tolerance against bacterial growth. In *W. somnifera*, it is extensively reported that MeJA triggers the expression of genes involved in phytosterol biosynthesis and those related to withanolides biosynthesis resulting in enhanced accumulation of withanolides and withanosides (Chaturvedi and Mishra 2012; Singh et al. 2015, 2017; Pandey et al. 2016). Consistent with these reports, the expression of both WsGT4 and WsGT6 was induced after 48 hours of exposure to MeJA, indicating their role in secondary metabolism (Figure S4). Furthermore, both exhibited the highest expression in leaves compared to other tissues, suggesting their involvement in leaf-specific secondary metabolism (Figure 3). In a similar manner, the expression of three sterol glycosyltransferases and a flavonoid GT reported earlier in *W. somnifera*, was found to be highest in leaves (Chaturvedi and Mishra, 2012; Kumar et al., 2013). Sequence analysis revealed that both WsGT4 and WsGT6 possess, at the C-terminal end of the protein, a distinctive feature of GTs involved in secondary metabolism: the conserved HCGWNS motif within PSPG box that is crucial for the enzyme’s glycosyltransferase functionality (Gachon et al. 2005) (Figure 2B). It was observed that WsGT4 showed 75.78% identity with *Lycium barbarum* UGT73A10, followed by 73.54% identity to NtUGT103 from *Nicotiana tabacum* (Noguchi et al. 2008; Yang et al. 2023). UGT73A10 is involved in secondary metabolism and exhibited regio-specific GT activity towards flavan-3-ols, and possessed a high degree of specificity for UDP-glucose (Noguchi et al. 2008). WsGT6 has 86.29% identity to *Solanum lycopersicum* SlUGT75C1 and 85.59% identity to NtGT2 from *Nicotiana tabacum* (Taguchi et al. 2003; Sun et al. 2017). The tomato SlUGT75C1 is an abscisic acid (ABA) uridine diphosphate glucosyltransferase involved in ABA-mediated fruit development and ripening, seed germination, and responses to drought stress in tomato (Sun et al. 2017).

GTs in plants are known to utilize either UDP-glucose or UDP-galactose or both as sugar donors (Jones and Vogt 2001; Bowles et al. 2006). GT assay with recombinant WsGT4 showed that it catalyzed the product formation with withanolide A, withanolide B, 12-deoxywithastramonolide, and withanone, but only in the presence of UDP-glucose, and not with UDP-galactose. Besides, WsGT4 formed a single product with 12-deoxywithastramonolide, and two different products with withanolide A, withanolide B, and withanone. In contrast, WsGT6 formed multiple products only with withaferin A, being able to do so in the presence of either UDP-glucose or UDP-galactose. This suggested that WsGT4 is cofactor-specific but can accept multiple substrates in its active site to form different glycowithanolides, whereas WsGT6 is substrate-specific but can accept both co-factors. Moreover, the formation of two different products with withanolide A, withanolide B, and withanone by WsGT4, and the formation of multiple products with withaferin A by WsGT6 suggests the glycosylation of respective substrates at different sites by WsGT4 and WsGT6 which is evident by peaks formed at different retention times (Figures 4A, B & C; 5A & B). A similar scenario has been reported in other plants; for instance, three UDP-glucose lignan GTs from *Isatis indigotica* catalyzed glycosylation at 4-hydroxyl group and 4′-hydroxyl group of lariciresinol (Tan et al. 2022), whereas in *Fallopia multiflora*, FmUGT1 and FmUGT2 glycosylated multiple sites of resveratrol, 2, 3, 5, 4’-tetrahydroxystilbene-2-O-β-D-glucoside, piceatannol, kaempferol, myricetin, curcumin, and phloretin (Cai et al. 2022). Plant GTs characterized previously have been known to act on a variety of substrates, including phenylpropanoids, alkaloids, terpenoids, and polyketides (He et al. 2022). Similarly, earlier studies in *W. somnifera* have identified few GTs that act on sterols and flavonoids (Sharma et al. 2007; Kumar et al. 2013). Thus, the present study is the first report of GTs from *W. somnifera* that specifically act on withanolide substrates.

In previous studies, VIGS and transient overexpression approaches have been effectively employed to investigate the *in planta* function of genes related to the biosynthesis and regulation of withanolides (Singh et al. 2015, 2017; Shilpashree et al. 2022). As biochemical studies indicated the role of WsGT4 and WsGT6 in withanosides formation, VIGS and transient overexpression experiments were employed to further assess their *in planta* role in withanosides biosynthesis in *W. somnifera*. Consequently, specific silencing and overexpression of *WsGT4* and *WsGT6* resulted in drastic down-regulation and enhancement of their transcripts in VIGS and overexpressing samples, respectively. The level of silencing and overexpression observed in *WsGT4* and *WsGT6* was similar to the degree achieved in previous studies by our group and by other research groups targeting genes in *W. somnifera* (Singh et al. 2015, 2017; Agarwal et al. 2018; Shilpashree et al. 2022, 2024). The silencing of *WsGT4* resulted in significant increase of withanolide B, withanoside V levels along with non-significant but enhanced accumulation of withanolide A, withanoside IV, and 12-deoxywithastramonolide. In contrast, leaves overexpressing *WsGT4* showed an opposite trend in terms of accumulation of withanoside IV, withanoside V, and 12-deoxywithastramonolide. In the case of *WsGT6*, its silencing and overexpression led to respective enhancement and reduction in the accumulation of withanolide A and 12-deoxywithastramonolide. Interestingly, overexpression of *WsGT6* also led to an increase in withaferin A levels. These results underscore the involvement of *WsGT4* and *WsGT6* in the biosynthesis of withanosides and their indirect impact on the production of withanolides. Based on these results, it can be construed that WsGT4 could drive the carbon flux to yet unknown withanoside/s, thus depleting the precursors for other withanolides/sides, whereas WsGT6 could be involved in diverting the carbon flux away from withanolide A to a yet to be identified intermediate (Figure 9). Previous work on the RNAi-mediated suppression of the *WsSGTL1* gene led to a reduction in withanoside levels alongside a notable increase in withanolides and glycosylated sterols within the silenced plants. The transgenic RNAi lines exhibited heightened production of withanolides, notably withanolide A, and campesterol, stigmasterol, and β-sitosterol in their glycosylated states, along with decreased accumulation of withanosides such as withanoside V (Saema et al. 2015). Similarly, silencing of other *WsSGTL* members (*WsSGTL1, WsSGTL2,* and *WsSGTL4*) through aMIR-VIGS led to an increased accumulation of withanolides and phytosterols while reducing the content of glycowithanolides in the silenced lines (Singh et al. 2016).

**Figure 9:**
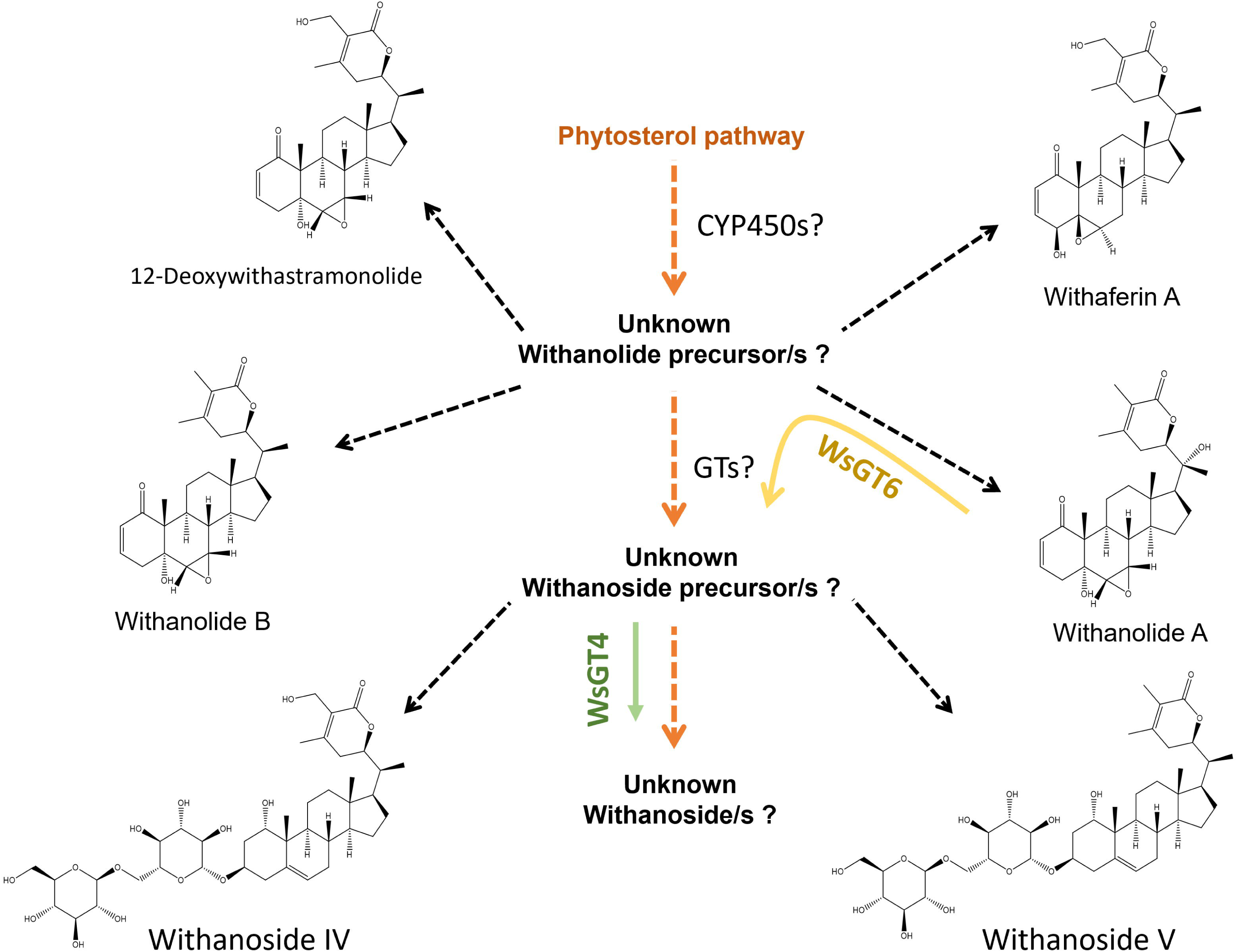
Proposed role of WsGT4 and WsGT6 in withanoside biosynthesis in *W. somnifera*. Based on metabolite data from silencing and transient overexpression, WsGT4 (green arrow) could drive the carbon flux to yet unknown withanoside/s, thus depleting the precursors for other withanolides/sides. WsGT6 (yellow arrow) could be involved in diverting the carbon flux away from withanolide A to a yet to be identified intermediate.

Glycosyltransferases play a crucial role in enhancing plant resistance to both biotic and abiotic stresses (Zhang et al. 2014). Specifically, UDP-GTs contribute to bolstering resistance against these stresses by modulating the activities of glycosylated hormones and secondary metabolites within plants (O’Donnell et al. 1998). In *W. somnifera*, increased levels of sterol glycosyltransferase expression were noted to confer resistance against diverse environmental stresses (Chaturvedi and Mishra 2012; Mishra et al. 2013, 2017, 2021, 2022; Pandey et al. 2014; Saema et al. 2016; Singh et al. 2016). In agreement with these results, leaves from the *WsGT4* and *WsGT6* silenced plants exhibited pronounced disease symptoms and endured greater tissue damage compared to leaves from control plants (Figure S9A). Moreover, the bacterial growth assay revealed a substantial increase in the proliferation of *P. syringae* DC3000 in *WsGT4*-VIGS and *WsGT6*-VIGS samples. In contrast, overexpression of *WsGT4* and *WsGT6* resulted in increased resistance against bacterial proliferation in *W. somnifera* leaves that exhibited lower level of necrosis in comparison to the leaves of control plants (Figure S9B). Previously, Mishra et al.’s study showcased that elevating *WssgtL3.1* expression substantially improved immunity against *Pseudomonas syringae* in *A. thaliana* plants (Mishra et al. 2022). Thus, it is evident that besides their role in withanosides biosynthesis, *WsGT4* and *WsGT6* play a role in defence against bacteria, although the precise mechanism requires additional exploration. It is plausible that the products generated by these enzymes might contribute to fostering tolerance either through their antibacterial properties or by participating in signaling processes. In conclusion, this study has presented biochemical evidence of WsGT4 and WsGT6 utilizing specific withanolides as substrates to form withanosides. In addition, the study demonstrated the *in planta* role of *WsGT4* and *WsGT6* in withanosides biosynthesis and defence against a model bacterial pathogen. These characterized GTs hold significant potential for metabolic engineering in *W. somnifera* cell cultures or the entire plant, facilitating targeted enhancement of specific withanolides/withanosides and the development of plants tolerant to bacterial pathogens.

## Author contributions

P.A., A.K.N., and D.P. performed the experiments. P.A., A.K.N., D.P., A.P., and D.A.N. analyzed the data. D.A.N. conceived and coordinated the research. P.A., A.K.N., D.P., A.P., and D.A.N. wrote the manuscript.

## Acknowledgements

A.K.N., D.P., and A.P. are recipients of research fellowships from Indian Council of Medical Research (ICMR), Department of Biotechnology (DBT), and Council of Scientific and Industrial Research (CSIR), India respectively. The authors are thankful to the Director, CSIR-CIMAP, for the support throughout the study. This manuscript bears the institutional communication number CIMAP/PUB/ 2024/23. Authors declare no conflict of interest.

## Data availability statement

All the data generated or analyzed during the study are available from the corresponding author upon request.

