## Supplemental data for "Functional characterization of two glycosyltransferases from *Withania somnifera* illuminates their role in withanosides biosynthesis and defence against bacteria"

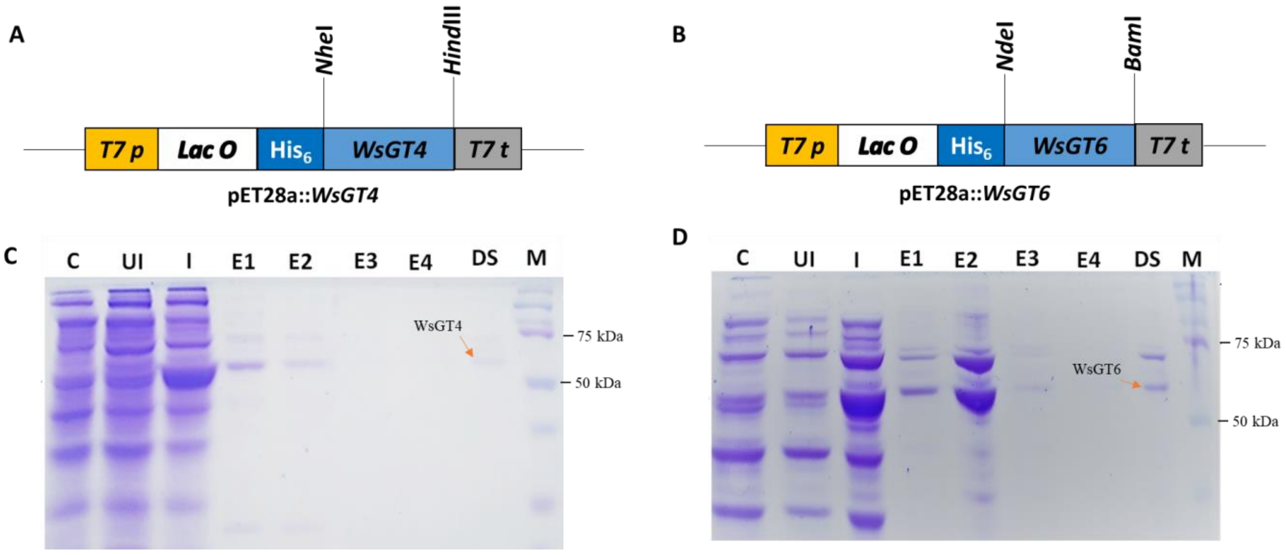


**Figure S1:** Purification of recombinant WsGT4 and WsGT6 proteins. **A)** Pictographic representation of pET28a::*WsGT4* and **B)** pET28a::*WsGT6* DNA constructs. **C)** SDS-PAGE showing total protein from crude cell lysate of empty vector control cells 'C', uninduced cells 'UI', induced cells 'I', and consequent eluted protein after Ni-affinity purification 'E1-E4' as well as pooled, purified and desalted protein 'DS' of WsGT4 expressing cells and **D)** that of WsGT6 expressing cells. Desalted proteins were quantified and used for assay.

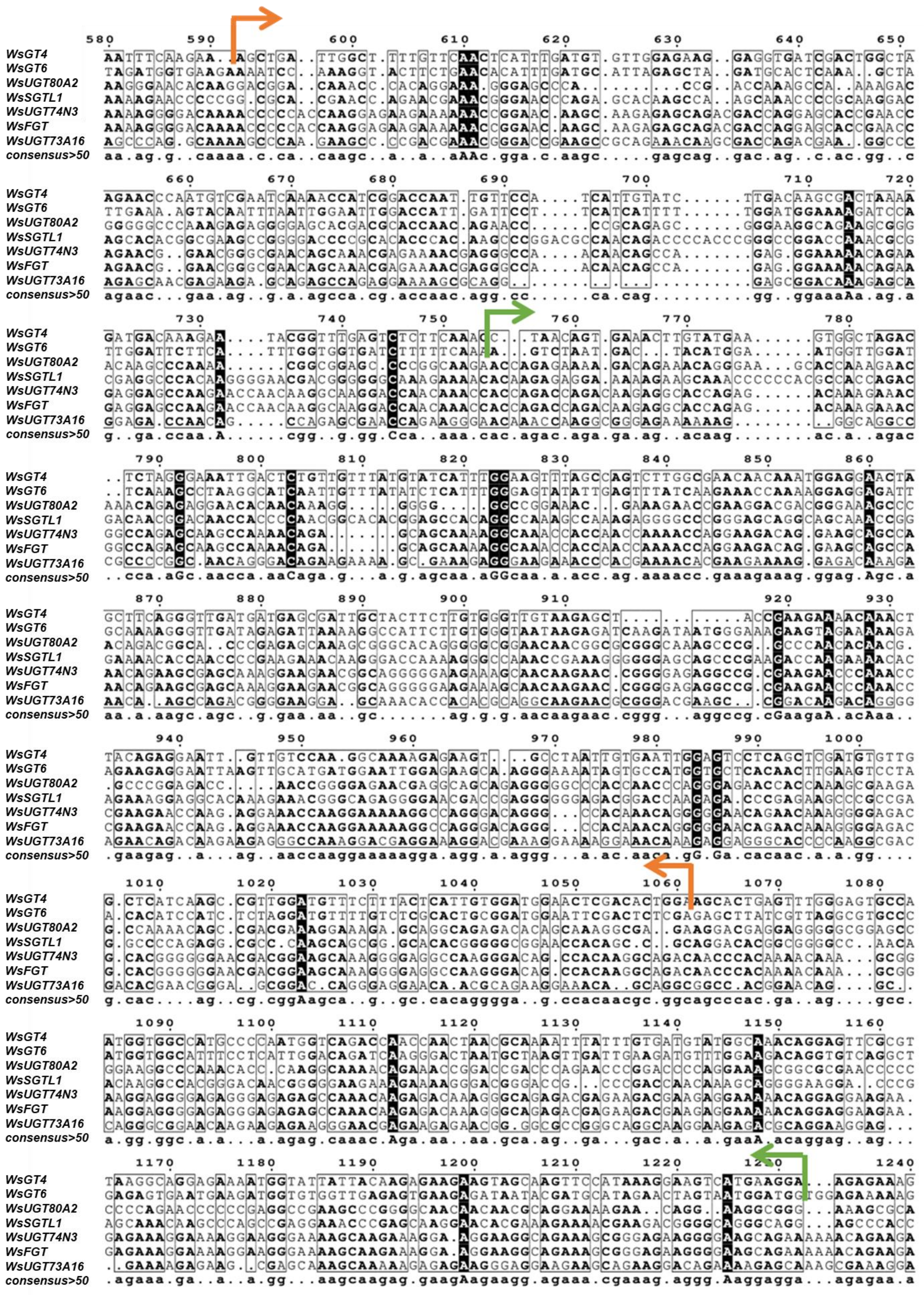

**Figure S2:** Multiple sequence alignment of WsGT4 and WsGT6 with other glycosyltransferases from *W. somnifera*. Sequence between orange and green arrows indicate the 471bp and 493bp region used for *WsGT4*-VIGS and *WsGT6*-VIGS respectively. The alignment shows very low consensus between the sequences in the region chosen for VIGS, greatly reducing the likelihood of off-target silencing. Accession numbers are as follows, *WsUGT80A2*: Z83833, *WsUGT74N3*: FJ560880, *WsUGT73A16*: FJ654696, *WsFGT*: FJ560880, *WsSGTL1*: DQ356887.

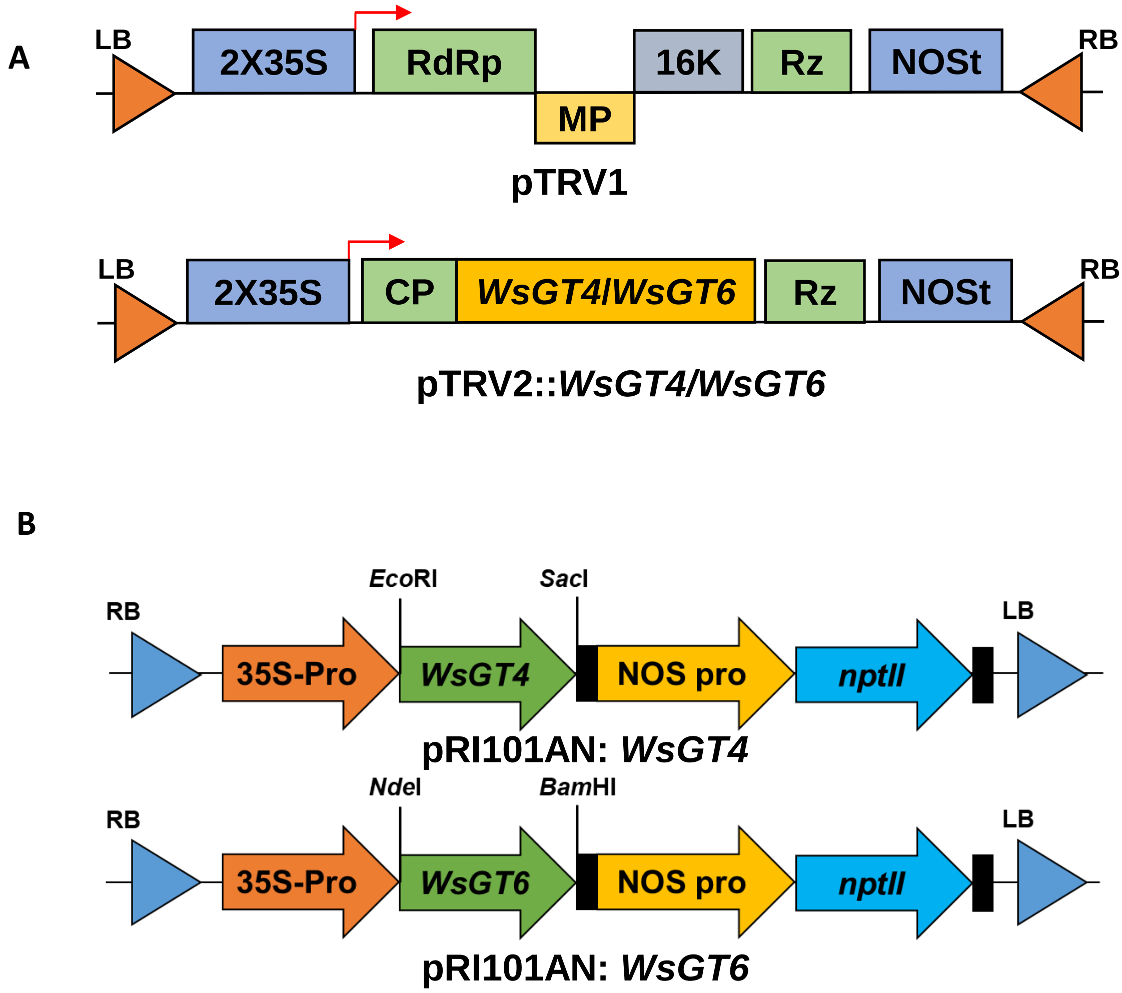


**Figure S3. A)** Vector map of pTRV1 and pTRV2::*WsGT4/WsGT6* constructs used in Virus-Induced Gene Silencing. **B)** Vector map of pRI101AN: *WsGT4/WsGT6* constructs used in transient overexpression study.

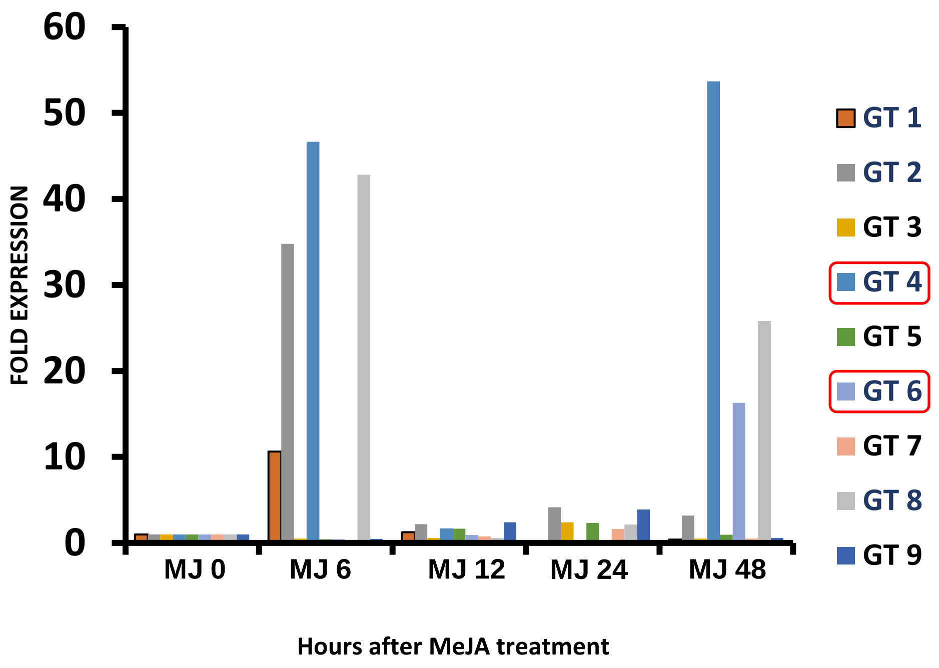


**Figure S4.** Effect of MeJA treatment on expression of candidate GTs. Leaves were treated with MeJA, tissues were collected at various time intervals 0 h, 6 h, 12 h, 24 h and 48 h post treatment. Total RNA was isolated and the specific expression of candidate GTs was analyzed through qRT-PCR. The relative expression of GTs is represented as fold change and that of 0 h was set to 1.

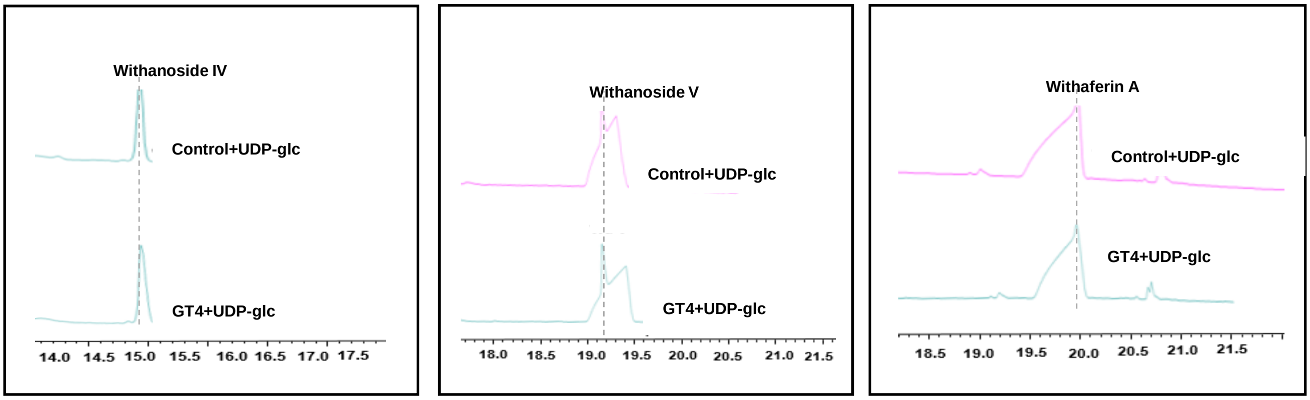


**Figure S5:** Biochemical assay using recombinant WsGT4 with withanolides and UDP-Glucose that did not form product.

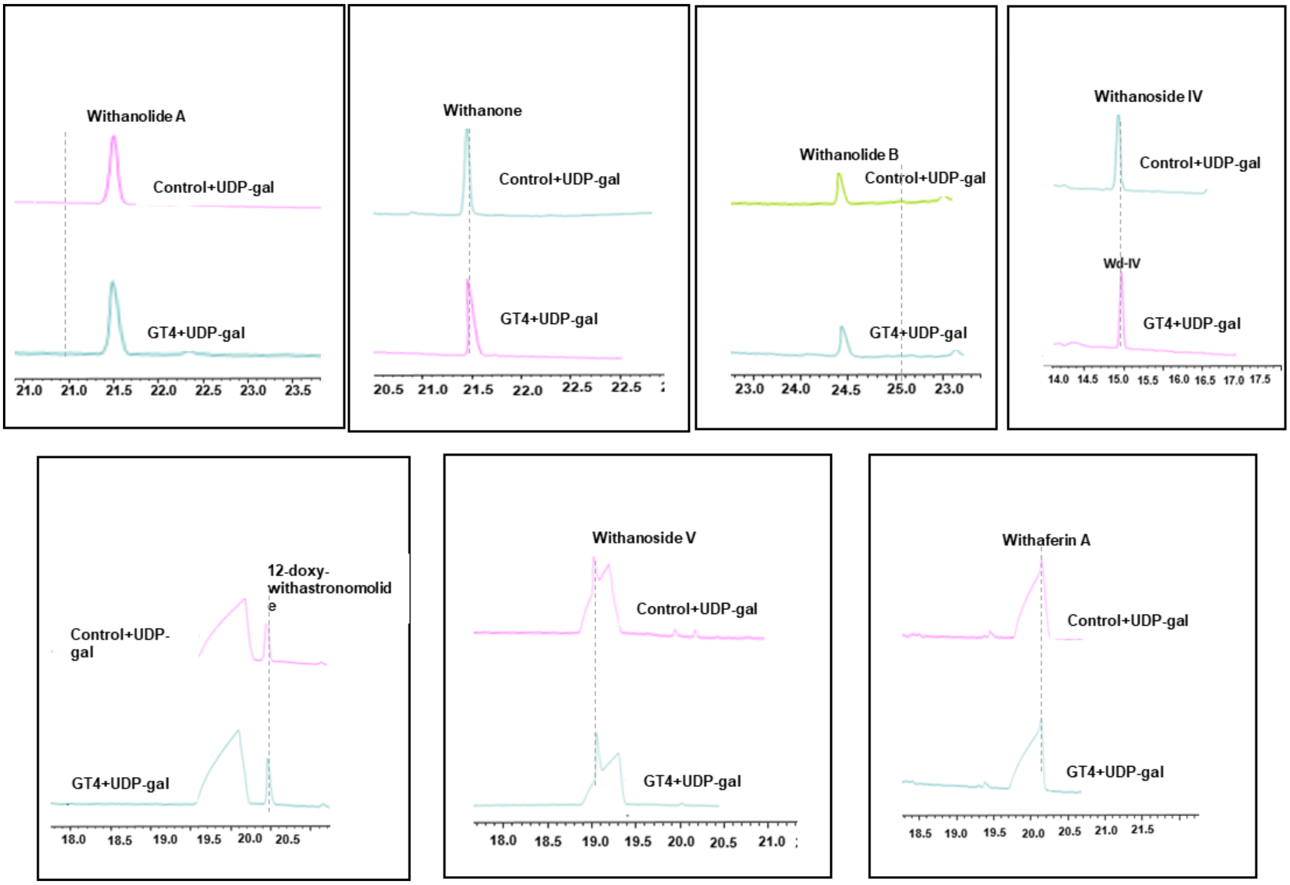


**Figure S6:** Biochemical assay using recombinant WsGT4 with withanolides and UDP-Galactose as a donor substrate that did not form product.

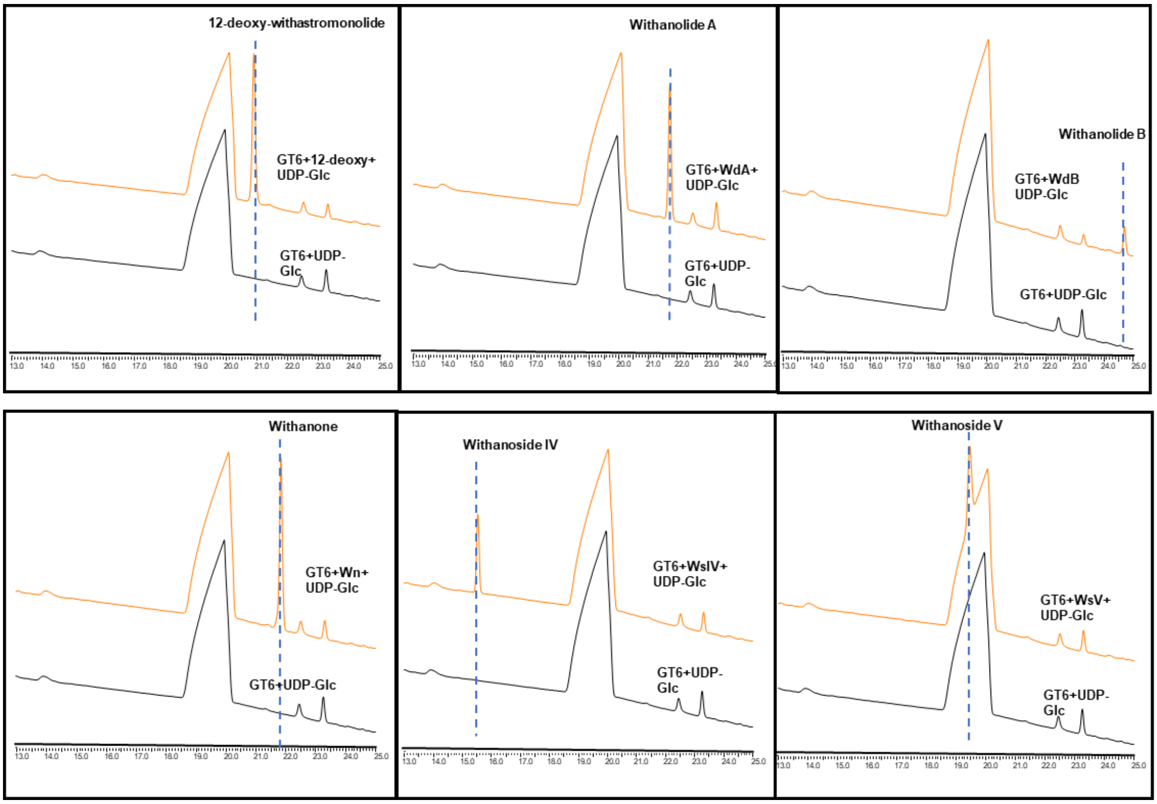


**Figure S7:** Biochemical assay using recombinant WsGT6 with withanolides and UDP-Galactose as a donor substrate that did not form product

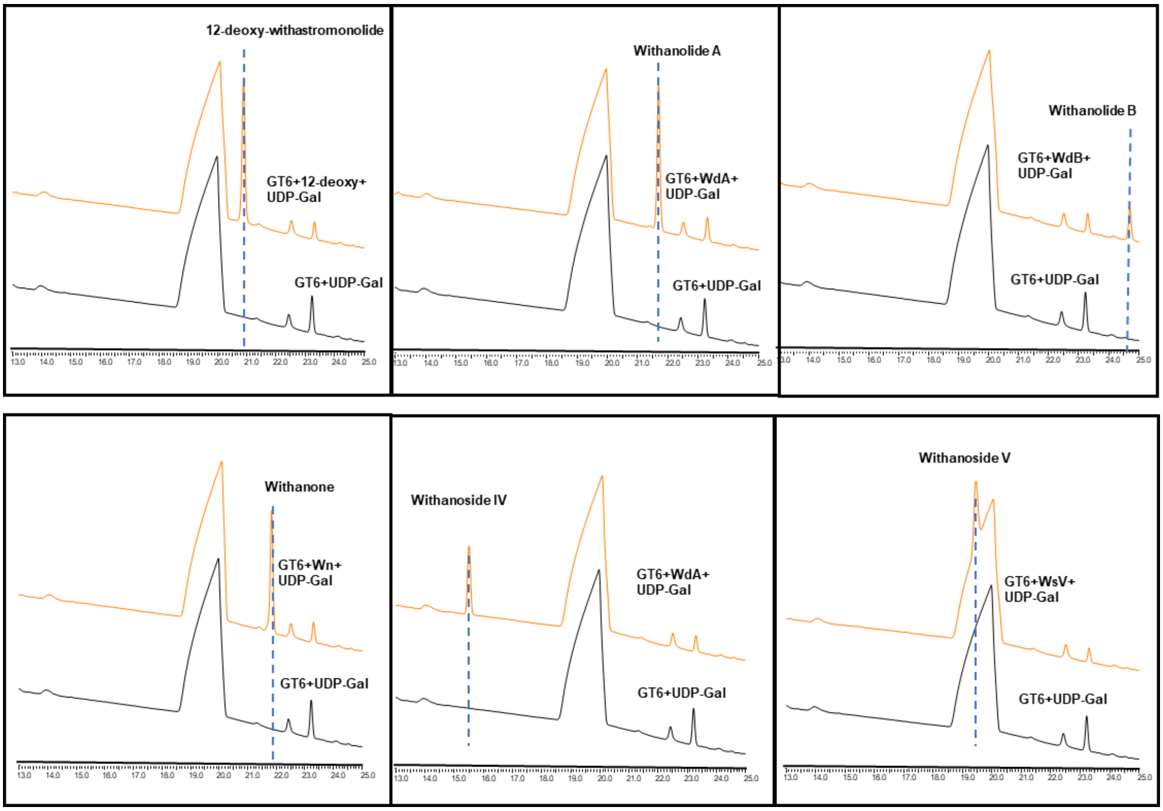


**Figure S8:** Biochemical assay using recombinant WsGT6 with withanolides and UDP-Glucose as a donor substrate that did not form product

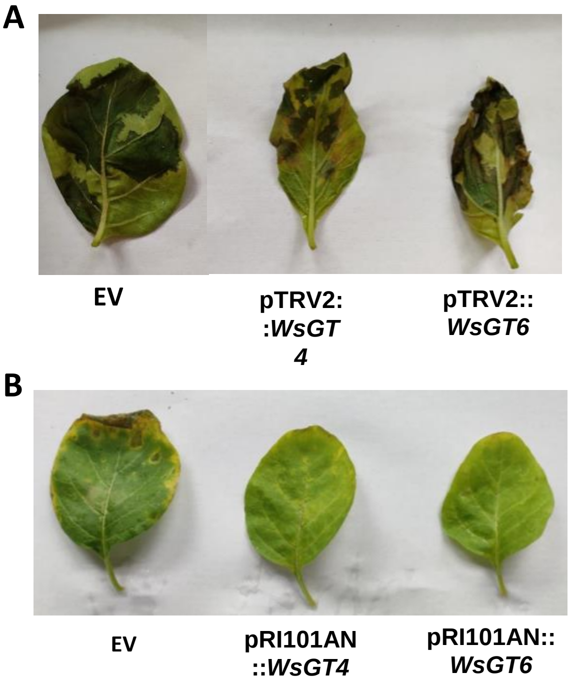


**Figure S9:** Phenotype of *W. somnifera* leaves infiltrated with *Agrobacterium* cultures harbouring different constructs and treated with *P. syringae*. **A)** Leaves infiltrated with VIGS constructs, pTRV2 empty vector (EV), pTRV2::*WsGT4* and pTRV2::*WsGT6*. **B)** Leaves infiltrated with overexpression constructs, pRI101AN empty vector control (EV), pRI101AN::*WsGT4* and pRI101AN::*WsGT6*.

**Table S1:** List of oligonucleotides used in the study.

| Primers | Sequence | Purpose |
|---|---|---|
| WsGT4-RT-F | GCTGATTGGCTTTTGTTCAACTC | qRT-PCR |
| WsGT4-RT-R | GGTCCGATGGTTTTGATTCG | qRT-PCR |
| WsGT6-RT-F | GAATTGGAGAAGCAAGGGAAAA | qRT-PCR |
| WsGT6-RT-R | CGAGACAAAACATCCTAGAGATGG | qRT-PCR |
| WsGT4-pET28a-F | GGCTAGCATGGAAAAACTACTCAACAAATCTCATG | pET28a cloning |
| WsGT4-pET28a-R | GAAGCTTTCAGTACATTACAATTTTGAGAAATTCTTCG | pET28a cloning |
| WsGT6-pET28a-F | GCATATGGCACAACCCCATGTACTCT | pET28a cloning |
| WsGT6-pET28a-R | GGGATCCTCAACAACCTTTGCCAACTT | pET28a cloning |
| WsGT4-pTRV2-F | TCTAGAGCTGATTGGCTTTTGTTCAACTC | pTRV2 cloning |
| WsGT4-pTRV2-R | CTCGAGTCCAGTGTCGAGTTCCATC | pTRV2 cloning |
| WsGT6-pTRV2-F | TCTAGAGTCTAATGACTACATGGAATGG | pTRV2 cloning |
| WsGT6-pTRV2-R | GGATCCCCATCCATTACTAGTTCTATGC | pTRV2 cloning |
| WsGT4- pRI101-AN-F | GAATTCATGGAAAAACTACTCAACAAATCTCATG | pRI101-AN cloning |
| WsGT4- pRI101-AN-R | GAGCTCTCAGTACATTACAATTTTGAGAAATTCTTCG | pRI101-AN cloning |
| WsGT6- pRI101-AN-F | GCATATGGCACAACCCCATGTACTCT | pRI101-AN cloning |
| WsGT6- pRI101-AN-R | GGGATCCTCAACAACCTTTGCCAACTT | pRI101-AN cloning |
